# Environmental enrichment promotes adaptive responding during tests of behavioral regulation in male heterogeneous stock rats

**DOI:** 10.1101/2023.06.30.547228

**Authors:** Keita Ishiwari, Christopher P. King, Connor D. Martin, Jordan A. Tripi, Anthony M. George, Alexander C. Lamparelli, Apurva Chitre, Oksana Polesskaya, Jerry B. Richards, Leah C. Solberg Woods, Amy Gancarz, Abraham A. Palmer, David M. Dietz, Suzanne H. Mitchell, Paul J. Meyer

## Abstract

Organisms must regulate their behavior flexibly in the face of environmental challenges. Failure can lead to a host of maladaptive behavioral traits associated with a range of neuropsychiatric disorders, including attention deficit hyperactivity disorder, autism, and substance use disorders. This maladaptive dysregulation of behavior is influenced by genetic and environmental factors. For example, environmental enrichment produces beneficial neurobehavioral effects in animal models of such disorders. The present study determined the effects of environmental enrichment on a range of measures related to behavioral regulation using a large cohort of male, outbred heterogeneous stock (HS) rats as subjects to mimic the genetic variability found in the human population. Subjects were reared from late adolescence onwards either in pairs in standard housing with minimal enrichment (n=200) or in groups of 16 in a highly enriched environment consisting of a large multi-level cage filled with toys, running wheels, and shelters (n=64). Rats were subjected to a battery of tests, including: (i) locomotor response to novelty, (iI) light reinforcement, (iii) social reinforcement, (iv) reaction time, (v) a patch-depletion foraging test, (vi) Pavlovian conditioned approach, (vii) conditioned reinforcement, and (viii) cocaine conditioned cue preference. Results indicated that rats housed in the enriched environment were able to filter out irrelevant stimuli more effectively and thereby regulate their behavior more efficiently than standard-housing rats. The dramatic impact of environmental enrichment suggests that behavioral studies using standard housing conditions may not generalize to more complex environments that may be more ethologically relevant.

## Introduction

Failure to regulate behavior to address environmental challenges is manifested in impaired inhibitory and attentional control, as well as aberrant responses to sensory and reward stimuli. Frequent failures of behavioral regulation are characteristics of a range of neuropsychiatric disorders, including attention deficit hyperactivity disorder (ADHD) autism spectrum disorders, schizophrenia, obesity, obsessive compulsive disorder, pathological gambling, and substance abuse ^e.g., 1–3^. While there is converging evidence for genetic bases of such dysregulatory behavioral traits ^e.g., 4, 5^, environmental factors are known to influence these and related traits as well ^e.g. 6–8^. For this reason, understanding gene-by-environment interactions in predicting susceptibility to various psychiatric disorders has been increasingly recognized by researchers ^9, 10^.

One way of studying the influence of environmental factors in preclinical research is by examining the behavioral effects of environmental enrichment (EE) ^11–13^. Experimentally, EE most often involves physical modifications of housing conditions such as the enhancement of cage space and the inclusion of objects and toys in the cage that allow animals to play, exercise, and explore. Social enrichment (i.e., group housing) is commonly included. Since the importance of EE was first described by Hebb ^14^, considerable preclinical evidence has indicated that EE produces a variety of beneficial neurobehavioral effects, including improved learning and memory and changes in neural structure ^e.g. 15–17^; although these effects may be moderated by genotype ^e.g., 18, 19^. Moreover, EE has been shown to exert beneficial effects in animal models of a wide variety of neurodegenerative and neuropsychiatric disorders ^20, 21^, including disorders associated with behavioral dysregulation such as ADHD, autism, schizophrenia, and substance abuse disorder ^e.g., 22–25^. It should be noted that in some scenarios environmental “enrichment” may have some negative consequences, such as an increase in anxiety and alcohol intake ^26, 27^. Thus, a better descriptor may be environmental complexity, although EE is used here to be consistent with recent literature.

The goal of the present study was to investigate the effects of EE on measures of behavioral regulation in a large cohort of genetically diverse heterogeneous stock (HS) rats, as part of the phenotyping component of large, multicenter genome-wide association study ^28–30^. Accordingly, we compared the performance of male HS rats that were group-housed in an environmentally complex cage from late adolescence onwards with those pair-housed in standard housing conditions, on tasks posited to measure different traits influenced by behavioral regulation. Specifically, to model sensation seeking we used two measures of stimulus reactivity: *locomotor response to novelty* and a *sensory reinforcement* test ^31^. We also used the *choice reaction-time, patch-depletion,* and *Pavlovian conditioned approach* tasks that measure different aspects of foraging behavior ^32, 33^; ^29^. Finally, we included tests of *social reinforcement* and *cocaine conditioned place preference*, to measure sociability and cocaine sensitivity, respectively ^34, 35^.

## Materials and Methods

### Subjects and Housing

Subjects were male heterogeneous stock (HS) rats shipped from the laboratory of Dr. Leah Solberg Woods, initially at the Medical College of Wisconsin (MCW: NMcwi:HS #2314009, RRID:RGD_2314009). Rats were shipped to the University of Buffalo in four batches (n = 66 / batch) at 4–5 weeks of age. Following quarantine (approximately one week), they were transported to the Clinical and Research Institute on Addictions. For each batch, on arrival at the Institute, 50 male rats were assigned to the standard housing (SH) condition, in which they were pair-housed in clear plastic cages (42 × 22 × 20 cm) lined with bedding (Aspen Shavings; Fig. 1B), conforming to the Guide for the Care and Use of Laboratory Animals ^36^. These rats were included as part of an ongoing genome-wide association study ^29, 30^, which is why there were more rats tested in this condition. The GWAS study ^29, 30^. The remaining 16 rats (from each of the 4 shipments, total n=64) were assigned to a single EE cage. Thus, 200 rats were assigned to SH and 64 to EE conditions. Because EE rats were not assigned to their conditions until after quarantine, the mean (± SEM) age of the SH rats and EE rats when they were placed into their housing assignments was 39.6 (± 0.6) days and 47.8 (± 0.4) days, respectively.

**Figure 1:**
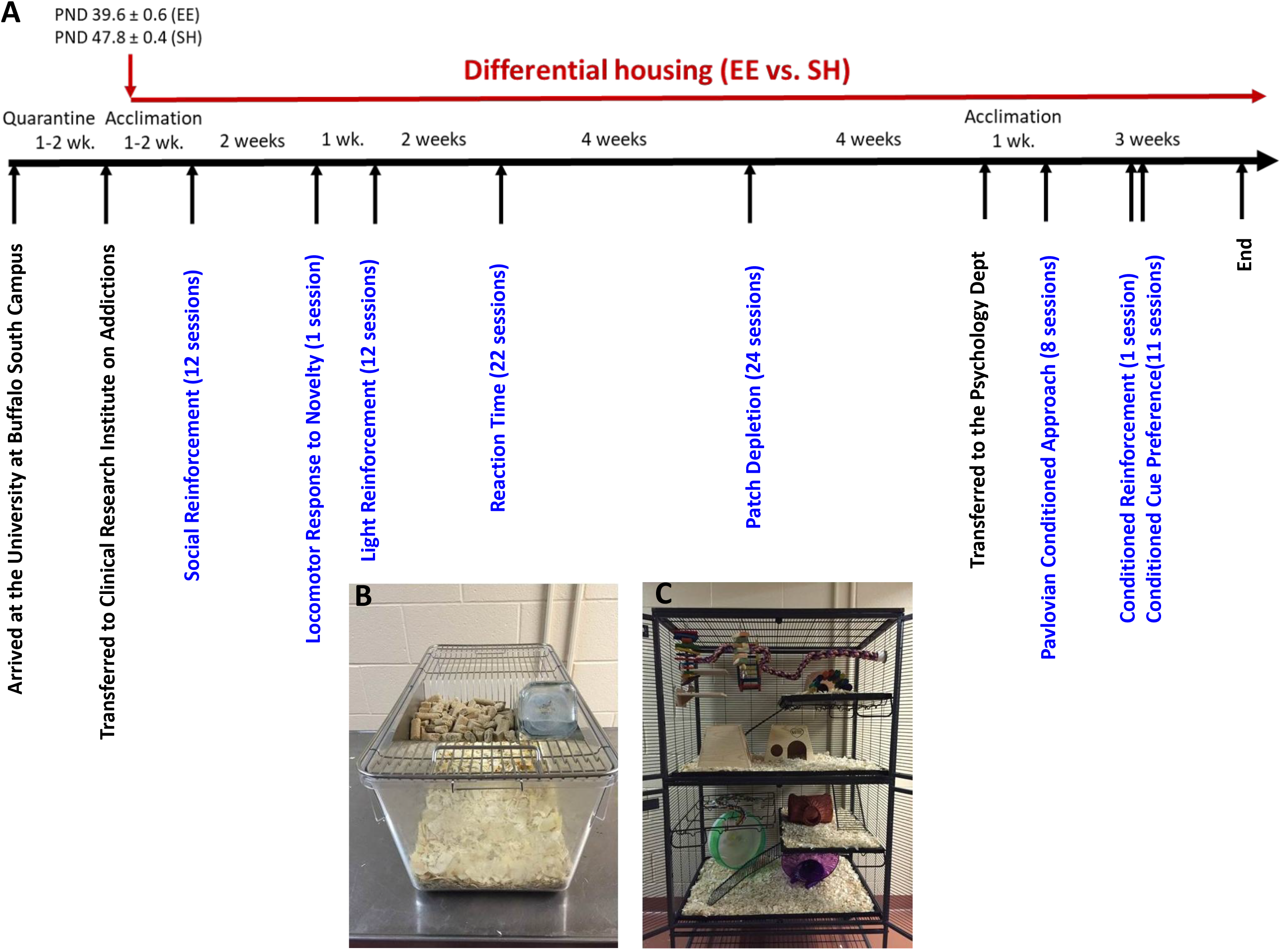
Experimental setup. (A) Approximate timeline of the study. Note that the order of the first two tasks was reversed for the third batch of 66 rats. Photographs show (A) the standard housing and (B) the environmentally enrichment cage.

As shown in Fig. 1C, the EE cage consisted of a large metal wire cage (90 × 60 × 120 cm; Doctors Forrest and Smith, Rhinelander, WI) with four levels connected by three ramps, each level containing a removable tray with bedding at the bottom. Approximately 15 plastic or wooden objects (e.g., huts, ladders, ropes, running wheels, bells, etc.) were placed inside the cage. The total area was 11,100 cm^2^, 12 times larger than the SH cage (924 cm^2^). Both SH and EE cages were in the same temperature (22 ± 1 °C) and humidity (55 %) controlled colony room on reverse light/dark cycle (lights on: 1900 to 0700 h). The bedding was replaced weekly.

Behavioral testing began after 1-2 weeks of acclimation to the reverse light/dark cycle. At the beginning of testing, the mean (± SEM) age of the EE rats was 49.8 (± 0.6) days, while that of the SH rats was 58.0 (± 0.4) days. Testing occurred 6 days a week during the dark phase of the light/dark cycle. Rats had *ad libitum* access to food (Teklad Laboratory Diet #8604, Envigo, Indianapolis, IN) except while in the testing chambers. Water was available *ad libitum* during all the behavioral tests except for the choice reaction time and patch-depletion tests, during which access to water was restricted to 30 min following testing. Rats were treated in compliance with the Guide for the Care and Use of Laboratory Animals, and the experiments were conducted in accordance with a protocol approved by the Institutional Animal Care and Use Committee (IACUC) at the University at Buffalo, the State University of New York.

### Behavioral testing procedures

Rats were tested on eight behavioral tasks (Fig. 1A): The first five tests were conducted at the Clinical and Research Institute on Addictions on the Medical Campus of the University at Buffalo, and then rats were transferred to the Department of Psychology for a one-week colony adaptation period followed by the last two tests. Below we briefly describe the apparatus, procedures and analysis pipeline for each behavioral test in the order rats completed them. Procedures are available on protocols.io with the exceptions of social reinforcement and cocaine cue preference: enriched housing (https://www.protocols.io/view/enriched-rat-housing-261gen2oyg47/v1) locomotor response to novelty (https://www.protocols.io/view/locomotor-response-to-novelty-cenntdde), reaction time (https://www.protocols.io/edit/reaction-time-testing-ceittcen), light reinforcement (https://www.protocols.io/edit/light-reinforcement-testing-cenptddn), patch depletion (https://www.protocols.io/view/delay-discounting-measured-using-a-sequential-patc-n92ldzqnnv5b/v1), and Pavlovian conditioned approach https://www.protocols.io/view/pavlovian-conditioned-approach-x54v9yjx4g3e/v2.

### Social reinforcement

#### Apparatus

Social behavior was measured using eight locally-constructed three-chamber apparatus ^34^. Briefly, the subject was placed in a circular center test chamber (diameter: 22.75 cm, height: 25.5 cm), which had three 2-inch observation ports with sliding doors opening into one of two circular stimulus chambers (diameter: 21.5 cm, height: 16.5 cm) located on the right and left sides of the test chamber or to the rear of the apparatus. Infrared photo sensors bisected the ports and detected snout pokes. Each port door was opened by operating a 24-volt rotating solenoid, which allowed physical contact between the subject and stimulus rats and the free passage of odor cues.

#### Procedure

A social stimulus (stimulus rat) was first placed into either the left or right stimulus chamber, counterbalanced across rats but fixed for each subject throughout social reinforcement testing. The stimulus rat was always a cage mate of the subject. During the 18-min test, the subject was placed into the center chamber, and the number of snout pokes into the three observation ports was recorded. Each door opened according to its own variable-interval (VI) 1-min schedule. Each rat was tested 3 days per week for a total of six sessions across two weeks, with each rat alternating as the subject or stimulus rat. Data from the last three tests for each rat were combined and used for analysis.

#### Analysis

The primary dependent measures were (1) the total number of responses in the social observation port, (2) the relative frequency of responses to the social observation port, obtained by dividing the number of responses to the social observation port by the sum of the responses to the social observation port and to the control observation port, and (3) interaction time, which was the amount of time the rat had its snout inside the social observation port when the door to the social stimulus chamber was open. The relative frequency of social responses indicates the reinforcing effectiveness of the social stimulus. For statistical analysis, the total numbers of responses from the last three sessions were analyzed using a three-way mixed factor ANOVA with housing condition (SH vs. EE) as a between-subjects factor and observation port (social, control) and time (six 3-min time epochs) as within-subjects factors. Relative frequency of social responses and interaction time when the social port was open was analyzed using an independent samples t-test to compare housing conditions.

### Locomotor response to novelty

#### Apparatus

Locomotor activity was recorded using eight infrared motion-sensor systems (Hamilton-Kinder Scientific, Poway, CA) as described in Gancarz et al. ^31^. Plastic cages (42 × 22 × 19 cm) were placed in sound- and light-attenuating enclosures with a ventilation fan. Infrared motion sensors were set at 5.5 and 15.5 cm above the cage floor on the outside of the cages. Lower lever sensors consisted of eight pairs along the long axis and five pairs along the short axis, each spaced 5.5 cm apart,. Upper level sensors were spaced 5.5 cm apart along the short axis only, and identified vertical rearing movements.

#### Procedure and Analysis

Each rat was placed into a novel, dark locomotor activity chamber for 18 min, and locomotor activity was recorded during a single session. The primary dependent measures were distance traveled, number of rears, and time spent in the center of the chamber. Distance traveled and rearing were divided into six 3-min time epochs (i.e., 0–3 min, 4–6 min, etc.) for analysis using a two-way mixed factor ANOVA with housing condition (SH vs. EE) as a between-subjects factor and time epoch as a within-subjects factor. Time spent in the center was analyzed using an independent samples t-test to compare housing conditions.

### Light reinforcement

#### Apparatus

Light reinforcement was assessed in 24 previously described operant chambers ^31, 32, 37^. Briefly, each test chamber had three snout poke holes located in the left, right and rear aluminum walls. Only pokes in the left and right holes were recorded using infrared photodetectors. Stimulus lights were located above each snout poke hole, and a house light was located in the ceiling of the test chamber.

#### Procedure

Rats were habituated to dark experimental chambers in six 18-min test sessions. During this pre-exposure phase, snout pokes had no programmed consequences but were recorded. Then there were six 18-min sessions in which rats were tested for operant responding for the light stimulus. During this light reinforcement phase, the test chambers were dark but a snout poke in the hole designated as “active” (left or right, counterbalanced) resulted in illumination of the ceiling stimulus light for 5 s according to a VI 1-min schedule of reinforcement. Snout pokes to the “inactive” alternative had no programmed consequences but were recorded.

#### Analysis

The primary dependent measures were (1) number of responses to the alternative that produced the light stimulus (active responding), (2) number of responses to the alternative that had no programmed effect (inactive responding), and (3) the relative frequency of active responding (defined as the number of active responses divided by the number of active and inactive responses).The relative frequency of active responses provides a measure of the reinforcing effectiveness for the response alternative that produced the visual stimulus ^38, 39^. Responding was analyzed using a three-way mixed factor ANOVA with housing condition (SH vs. EE) as a between-subjects factor and port (active vs. inactive) and session as within-subjects factors. Data on within-session changes in responding (numbers of active and inactive responses per 3-min time epoch) were also analyzed separately for the pre-exposure and light reinforcement phases using a three-way ANOVA with housing condition as a between-subjects factor and port and epoch as within-subjects factors.

### Reaction time

#### Apparatus

The reaction time task was conducted in 16 locally constructed experimental chambers as previously described ^32, 40, 41^. Briefly, the test panel had two water dispensers located on either side of a centrally located snout-poke hole. Stimulus lights were mounted above the two water dispensers and the center snout poke hole. Sonalert® tone generators were mounted above stimulus lights on the left (pure tone, 2.9 kHz) and right (pulsed tone, 1.9 kHz). Only the left tone generator and stimulus light were used. Snout pokes and head entries into the water dispensers were monitored with infrared detectors. Precise amounts of water were delivered to the water dispensers by syringe pumps (3.33 rpm PHM-100; Med Associates, St. Albans, VT).

#### Procedure

On a series of trials, a water-restricted rat could earn 30 μl of water by poking its snout into the center hole in the chamber and holding it there for a specific duration (“hold time”). The hold time was cumulative; for example, for a 1.6 s hold time, the rat could hold its snout in the hole for 0.8 s on two different occasions. A tone played while the rat made the snout poke response into the center hole. Once the hold time had elapsed, a stimulus light above the left hole (“imperative stimulus”) turned on to signal the availability of a water reinforcer in that hole. The rat had to remove its snout from the center hole and insert it into the dispenser within 3 s to earn the water reinforcer (a “correct response”). If the rat did not make a response within 3 s, the trial ended, and the trial was recorded as an omission. If the rat made an “incorrect response” (any response to the wrong dispenser), the trial ended without reinforcement. A “false alarm” was defined as a withdrawal from the center hole followed by a snout poke response into the left water dispenser hole prior to the presentation of the imperative stimulus.

Sessions lasted for 18 min, and training occurred 6 days a week. Initial training consisted of eight sessions requiring only a brief snout poke (minimal hold time of 0.1 s), into the center hole to initiate the trial. During Sessions 9–15, the hold time was systematically increased to a final value of 1.6 s, and data from Sessions 20–22 were averaged for the analysis.

#### Analysis

The primary measures were the number of trials completed and the mean reaction time. Trials completed provides a measure of attention because the completion of each trial requires effortful attention while waiting for the imperative stimulus. Mean reaction time was defined as the time elapsed from onset of the imperative stimulus to the rat’s head entering the water dispenser below the stimulus light for each rat. The number of false alarms was divided by the total number of trials initiated to provide an estimate of a proportion of premature responses that was not biased by the number of trials completed (false alarms per opportunity). Data on the number of trials completed were divided into six 3-min time epochs and were analyzed using a two-way ANOVA with housing condition (SH vs. EE) as a between-subjects factor and epoch as a within-subject factor. Mean reaction time and the number of false alarms per opportunity were analyzed using independent samples t-tests to evaluate the effect of housing condition.

### Patch Depletion

#### Apparatus

These chambers were the same as the ones used for the light reinforcement task, but also included water dispensers located inside of the left and center snout poke holes by syringe pumps (3.33 rpm PHM-100; Med Associates).

#### Procedure

The sequential patch depletion is a foraging task procedure described previously ^32, 42^, Figure 6A, which is loosely based on the adjusting amount procedure ^43^ and is designed to mimic naturally occurring choice problems confronting animals while foraging in an environment in which continued foraging at a specific location (patch) depletes the patch and decreases net rate of resource intake. A travel time delay is imposed when the animal moves from a depleted patch to a new patch. If the delay is long, optimal foraging theory predicts that the animal will deplete patches to lower levels before leaving ^44, 45^.

In this task, water-restricted rats drank water at the left and center water dispensers (patches). Rats received successively smaller amounts of water every 4 s by remaining at the same dispenser: the amount of water was initially 150 µl and then decreased by 20% after each delivery from the same dispenser (e.g., 150 µl at 0 s, 120 µl at 4 s, 96 µl at 8 s, 77 µl at 12 s, etc.). Rats could leave this patch by making a snout poke into an alternative dispenser at any time. This action would turn off the light above the original dispenser, and start a timer during which water was not available at either dispenser (“changeover delay”), signaled by a pulsed 1.9 kHz tone. At the end of the changeover delay, the light above the alternative dispenser would turn on and a snout poke would yield 150 µl water, diminishing according to the same pattern as described for the original dispenser. At any time, the rat could make a snout poke in the original dispenser, and the same changeover contingencies would be applied. Importantly, when during the 0-s delay condition, the most efficient strategy is to change patches after each water delivery. Each session lasted for 10 min or until the rat consumed a cumulative total 5,000 µl of water, whichever occurred first. Different changeover delays of 0, 6, 12, 18 and 24 s were imposed during different sessions, but were consistent within each session. Delays were tested in the following sequence in each week; 0, 0, 6, 12, 18, and 24 s. This cycle was repeated four times for a total of 24 test sessions.

#### Analysis

Data from the first day of the week with a 0-s delay were not included in the final statistical analysis. The primary dependent measures were the volume of water at which rats left a patch (“indifference point”) at each delay and the water consumption rate (in µl/min) at each delay. The indifference point indicates an equivalency between 150 µl reinforcer available following the changeover delay and the reinforcing effectiveness (or subjective value) of the final reinforcer earned from the depleting patch. Data on the indifference point and water consumption rate at each delay were both analyzed using a two-way ANOVA with housing condition (SH vs. EE) as a between-subjects factor and delay (0, 6, 12, 18 and 24 s) as a within-subjects factor.

### Pavlovian conditioned approach and conditioned reinforcement

#### Apparatus

This approach was based on previous studies in our laboratory and others ^29, 46^. Rats were tested in 16 conditioning chambers (21 × 24 × 29 cm; Med Associates) equipped with LED-illuminated retractable levers (2 cm long, 6 cm above floor) located on either the left or right side (counterbalanced) of a food cup (3 cm above a stainless steel grid floor). Banana-flavored food pellets (45 mg, BioServ, #F0059, Frenchtown, NJ) were delivered into the cup by an automatic dispenser. The food cup was equipped with an infrared photobeam that detected head entries. An illuminated red house light was located high (27 cm) on the opposite wall. For the conditioned reinforcement task, the food-cup was removed and the retractable lever was moved to the center of the wall in its place. Two snout-poke holes with photobeam detectors were located on the left and right side of the lever (counterbalanced across rats). All operant test chambers were housed in light and sound attenuating enclosures and controlled by computers connected to a Med Associates interface.

#### Procedure

On the 2 days prior to testing, all rats were fed 25 banana-flavored food pellets in their home cages. On the next day, a food cup training session occurred during which 25 pellets were dispensed into the food cup on a variable-time (VT) 30 s (range: 1-60 s) schedule of reinforcement, and food cup entries were recorded. Throughout the training session, the lever was retracted, and the red house light remained on. Starting on the following day, rats underwent five daily sessions of Pavlovian conditioned approach, each lasting 37.5 min on average. Each session consisted of 25 trials in which the 8-s illuminated lever presentation (conditioned stimulus: CS) was immediately followed by the retraction of the lever, extinguishing of the light, and delivery of a food pellet (unconditioned stimulus: US) into the food cup. Successive trials were separated according to a VT 90 s (30-150s) schedule. Food cup entries were recorded throughout the session during both the lever presentation and the inter-trial interval.

One day after the last Pavlovian conditioned approach session, we assessed the ability of the food-associated lever (CS) to act as a conditioned reinforcer when rats acquired a new instrumental response (snout poking). The conditioned reinforcement test was conducted in the same chamber, but the center food-cup was removed and replaced with the lever CS. On both the left and right side of the retracted lever-CS were two snout-poke ports, one active and one inactive. All other aspects of the testing environment were identical to the previous days. Rats were placed in the operant chamber for 40 min. Entry into the “active” snout-poke hole resulted in deployment of the lever into the chamber for 3 s according to a fixed-ratio (FR) 1 schedule, during which lever deflections were recorded, but additional entries into that snout-poke hole had no effects. Entries into the “inactive” hole had no programmed consequences. The numbers of entries into either hole, lever presentations earned, and lever contacts were recorded. Data on the number of snout pokes were divided into eight 5-min time epochs.

#### Analysis

For Pavlovian conditioned approach training, the numbers of lever contacts and food cup entries during the lever presentation and the inter-trial-intervals were analyzed using two-way ANOVAs with housing condition (SH vs. EE) as a between-subjects factor and session (5) as a within-subject factor. For conditioned reinforcement, active and inactive snout-poke responses were analyzed separately using two-way ANOVAs with housing condition as a between-subjects factor and epoch (8) as a within-subject factor. Data on the number of lever contacts per lever presentation were analyzed using an independent samples t-test for each housing condition.

### Cocaine conditioned cue preference

#### Apparatus

The conditioned cue preference task was conducted in 16 black acrylic chambers (47 × 19 × 30 cm) with smooth black matte floors located inside custom-made light-proof sound attenuating shells based on Meyer et al. ^29, 47^. Black spray-painted “grid” or “hole” textured floors were placed on top of the matte floor during conditioning. All sessions were videotaped using infrared cameras connected to a 16-channel DVR (Swann Communications, Santa Fe Springs, CA).

#### Procedure

On the first day (habituation), rats were injected with 0.9 % saline (1 ml/kg, i.p.) and placed into the chamber with only the smooth matte floor for 30 min in order to acclimate. On the following day, animals underwent a test to measure any preexisting floor bias (“pre-test”). During this test, they were injected with saline prior to being placed into their chamber containing “hole and “grid” floor halves (randomly assigned to the right and left). The least preferred floor (i.e., the floor the subject spent the least amount of time on) was assigned as the cocaine-paired floor for each individual. Rats were randomly assigned to either saline or cocaine conditions for the first day of conditioning. Conditioning lasted eight days in which rats were injected with cocaine (10 mg/kg) or saline i.p. on alternating days immediately before being placed into the dark chamber. Chambers contained only one floor type on these days based on drug pairing. Each set of a cocaine-paired and saline-paired day was termed a trial to yield four conditioning trials. Afterwards, rats underwent a post-conditioning test when they received a saline injection and were placed into the dark chamber with both floor types (configured the same as during the floor bias test). Each session lasted 30 min. Time spent on each floor type was recorded during the pre-test and the post-test.

#### Analysis

The primary dependent measure was time spent on the cocaine-paired floor type. Locomotor activity was also recorded on all test days. Data on saline- and cocaine-induced locomotor activity (distance traveled) during the conditioning trials were analyzed using a three-way ANOVA with housing condition (SH vs. EE) as a between-subjects factor and drug (cocaine, saline) and conditioning trials (4) as within-subjects factors. Data on time spent on the cocaine-paired floor were analyzed using a two-way ANOVA with housing condition as a between-subjects factor and test (pre- vs. post-test) as a within-subjects factor. Data comparing the change in time spent on the cocaine-paired floor between the housing conditions were analyzed using an independent samples t-test.

### Data collection and statistics

For most tasks, MED-PC IV software (Version 4.2, Med Associates) was used to apply the experimental contingencies and to collect data. Cocaine cue preference data (locomotion and side preference) was analyzed using TopScan video tracking software (CleverSys, Reston, VA). In the statistical analyses for all the tests, when the sphericity assumption was violated in conducting repeated-measures ANOVAs, a multivariate test (Wilks’ Ʌ) was used. Significant F values were followed by *post hoc* comparisons with Bonferroni corrections (0.05/number of comparisons). The level of significance was set at *p* < 0.05. Effect sizes were calculated using partial eta squared (η_p_^2^), Glass’s δ, or Cohen’s *d* as appropriate. Statistical analyses were carried out using SPSS Statistics (vers. 27, IBM, Armonk, NY) or Statistica (vers. 13, TIBCO, Palo Alto, CA).

### Drugs

Cocaine solution used for the conditioning cue preference test was prepared by dissolving cocaine HCl (National Institute of Drug Abuse, Bethesda, MD) into 0.9 % sterile saline at a concentration of 10 mg/ml to be administered at 10 mg/kg intraperitoneally (i.p.).

## Results

The direction and effect sizes of all tests conducted are presented in Table 1. The text below contains more specific results, which are also presented in the figures.

**Table 1.** Summary of major results. The major effects of housing are shown (n.s. = not significant); more detailed analyses are presented in the results. Asterisks indicate statistical trends with p<0.10 for goal-tracking (EE>SH) and locomotion during cocaine cue preference (SH>EE).

| **<u>Test</u>** | **<u>Major Measures</u>** | **<u>Effect</u>** | **<u>Effect Size</u>** |
| --- | --- | --- | --- |
| Social Reinforcement | Total Responses | SH>EE | $\eta_p^2 = 0.225$ |
|  | Relative Responses | n.s. |  |
| | Interaction Time | EE>SH | Glass's $\delta = 0.359$ |
| Locomotor Response to Novelty | Distance | SH>EE | $\eta_p^2 = 0.057$ |
| | Rearing | SH>EE | $\eta_p^2 = 0.207$ |
|  | Center Time | n.s. |  |
| Light Reinforcement | Total Responses | SH>EE | $\eta_p^2 = 0.364$ |
| | Relative Responses | SH>EE | $\eta_p^2 = 0.063$ |
| Reaction Time Task | Reaction Time | SH>EE | Cohen's $d = 0.496$ |
| | Trials Completed | EE>SH | $\eta_p^2 = 0.314$ |
| | Omissions | SH>EE | Glass's $\delta = 0.881$ |
| | False Alarms | SH>EE | Glass's $\delta = 0.744$ |
| Patch Depletion | Discounting Slope | EE>SH | $\eta_p^2 = 0.061$ |
| | Indifference Point | SH>EE | $\eta_p^2 = 0.232$ |
| | Water Consumed | EE>SH | $\eta_p^2 = 0.146$ |
| Pavlovian Conditioned Approach | Sign-tracking | EE>SH | $\eta_p^2 = 0.104$ |
|  | Goal-tracking | n.s.* |  |
| | Cond. Reinforcement | EE>SH | $\eta_p^2 = 0.239$ |
| Cocaine Cue Preference | Preference | n.s. |  |
|  | Locomotion | n.s.* |  |

### Social reinforcement

Fig. 2A shows the average number of snout poke responses into the social and control ports across the six time epochs averaged across the last three sessions of the social reinforcement test. The EE rats made fewer responses than the SH rats overall [main effect of housing on the number of responses: F(1, 262) = 76.24, *p* < 0.001, η_p_^2^ = 0.225]. Both groups made more responses into the social port than into the control port [main effect of port: F(1, 262) = 74.19, *p* < 001, η_p_^2^ = 0.221]. However, the difference in responding between the social and control ports was smaller for the EE rats than for the SH rats [housing × port interaction: F(1, 262) = 20.27, *p* < 0.001, η_p_^2^ = 0.072]: *post hoc* tests done separately for each group indicated that the EE rats responded significantly more to the social port only in Epochs 1, 5, and 6 (*p* < 0.0083 = 0.05/6), while SH rats responded significantly more to the social port than to the control port in all six epochs. However, as shown in Fig. 2B, there was a trend toward greater frequency of social response in the SH rats that did not reach significance [t(262) = 1.88, *p* = 0.061]. As for within-session changes in responding, there were a significant main effect of time epoch on responding [Wilk’s Ʌ = 0.34, F(5, 258) = 99.56, *p* < 001, η_p_^2^ = 0.659] as well as a significant interaction between port and time epoch [Wilk’s Ʌ = 0.90, F(5, 258) = 5.53, *p* < 0.001, η_p_^2^ = 0.097], indicating that overall responding declined within the session at different rates between the two ports. However, there was no significant interaction between housing type and epoch [Wilk’s Ʌ = 0.98, F(5, 258) = 1.15, *p* = 0.333], indicating that responding by the rats in EE and SH groups declined at similar rates. No significant three-way housing × port × epoch interaction was found [Wilk’s Ʌ = 0.99, F(5, 258) = 0.80, *p* = 0.553]. Finally, as shown in Fig. 2C, EE rats spent more time on average with their snout in the social observation port when the sliding door was open than the SH rats [t(63.37) = 2.87, *p* < 0.01, Glass’s *δ* = 0.359], indicating more social interaction time.

**Figure 2:**
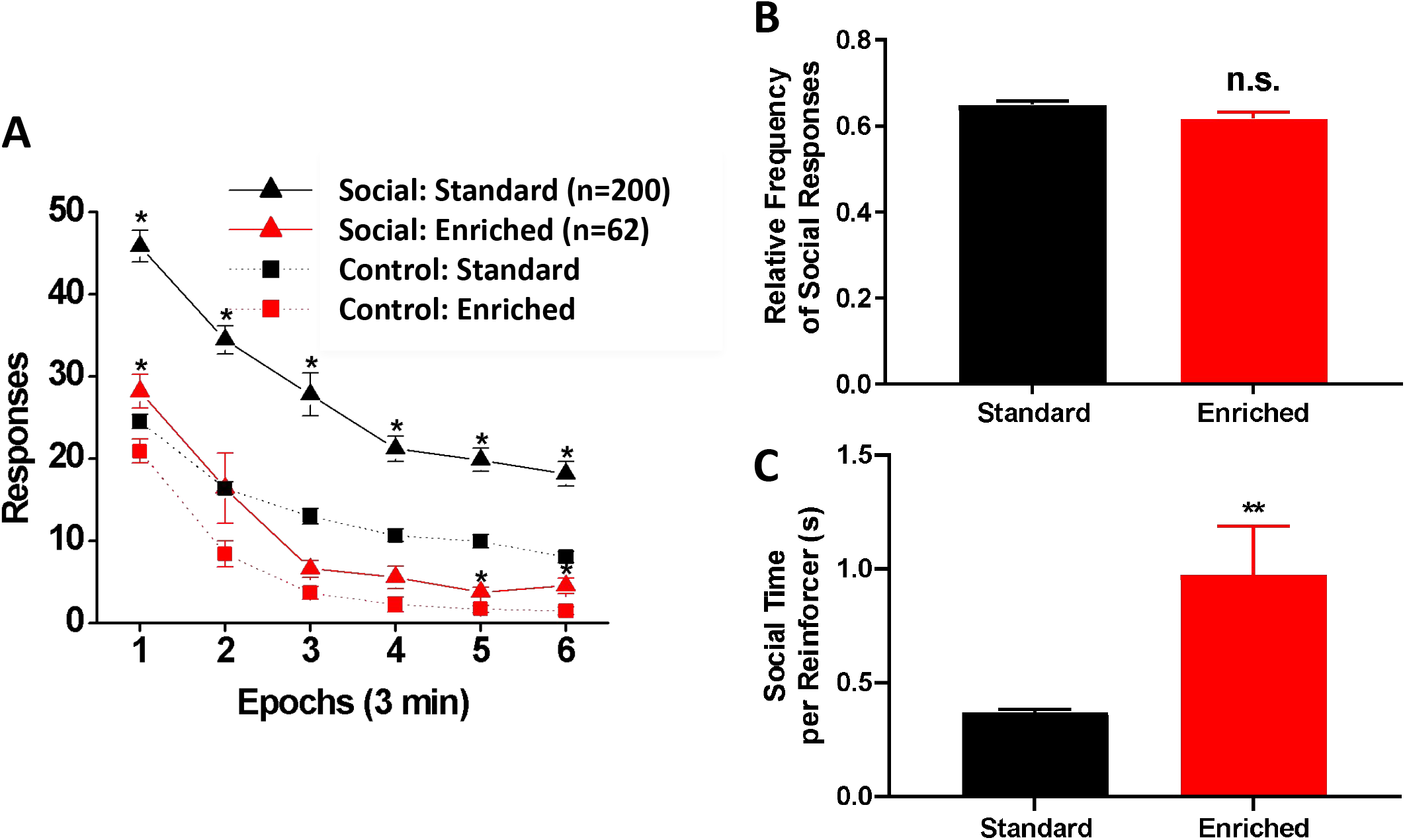
Social reinforcement test. (A) Within-session performance during social reinforcement testing in SH (black: n = 200) and EE (red: n = 64) rats. Average numbers of snout poke responses into the social (triangle) and opposite control (square) ports made by the SH and EE rats from the last three social reinforcement sessions are shown. Data are means (± SEM) across six time epochs during the session. * *p* < 0.0083 vs. opposite control. (B) The mean (± SEM) relative frequency of social responses from the last three sessions. While the SH rats showed a higher preference for the social stimulus than the EE rats, the difference did not reach statistical significance (*p* = 0.0613). (C) The mean (± SEM) time (in seconds) the rat spent with its snout in the social observation port when the sliding door was open. When given access, the EE rats spent more time contacting the social stimulus than the SH rats. ** *p* < 0.01.

### Locomotor response to a novel environment

Two EE rats escaped from the test chamber during the locomotor test session and were excluded from the analyses. As shown in Fig. 3A, the distance traveled in the novel locomotor chamber declined sharply across the 3-min epochs during the session for the rats in both housing conditions [main effect of time epoch: Wilk’s Ʌ = 0.104, F(5, 256) = 441.83, *p* < 0.001, η_p_^2^ = 0.896]. However, the EE rats traveled less than the SH rats [main effect of housing: F(1, 260) = 15.73, *p* < 0.001, η_p_^2^ = 0.057]. Moreover, the EE rats habituated to the novel locomotor chamber more quickly than the SH rats [housing × time epoch interaction: Wilk’s Ʌ = 0.658, F(5, 256) = 26.64, *p* < 0.001, η_p_^2^ = 0.342]. A similar pattern of results was obtained for rearing (Fig. 3B). There was a significant interaction between housing and time epoch [Wilk’s Ʌ = 0.852, F(5, 256) = 8.93, *p* < 0.001, η_p_^2^ = 0.148] as well as significant main effects of housing [F(1, 260) = 67.75, *p* < 0.001, η_p_^2^ = 0.207] and time epoch [Wilk’s Ʌ = 0.131, F(5, 256) = 338.73, *p* < 0.001, η_p_^2^ = 0.869]. Finally, there was no significant difference in the time spent in the center of the locomotor chamber [t(260) = 0.24, *p* = 0.808] (Fig. 3C). Thus, the present results demonstrated that, while rats in both housing conditions displayed clear declines in locomotor activity within the test session, the EE rats showed a smaller locomotor response in a novel environment and habituated to the novel environment faster relative to the SH rats.

**Figure 3:**
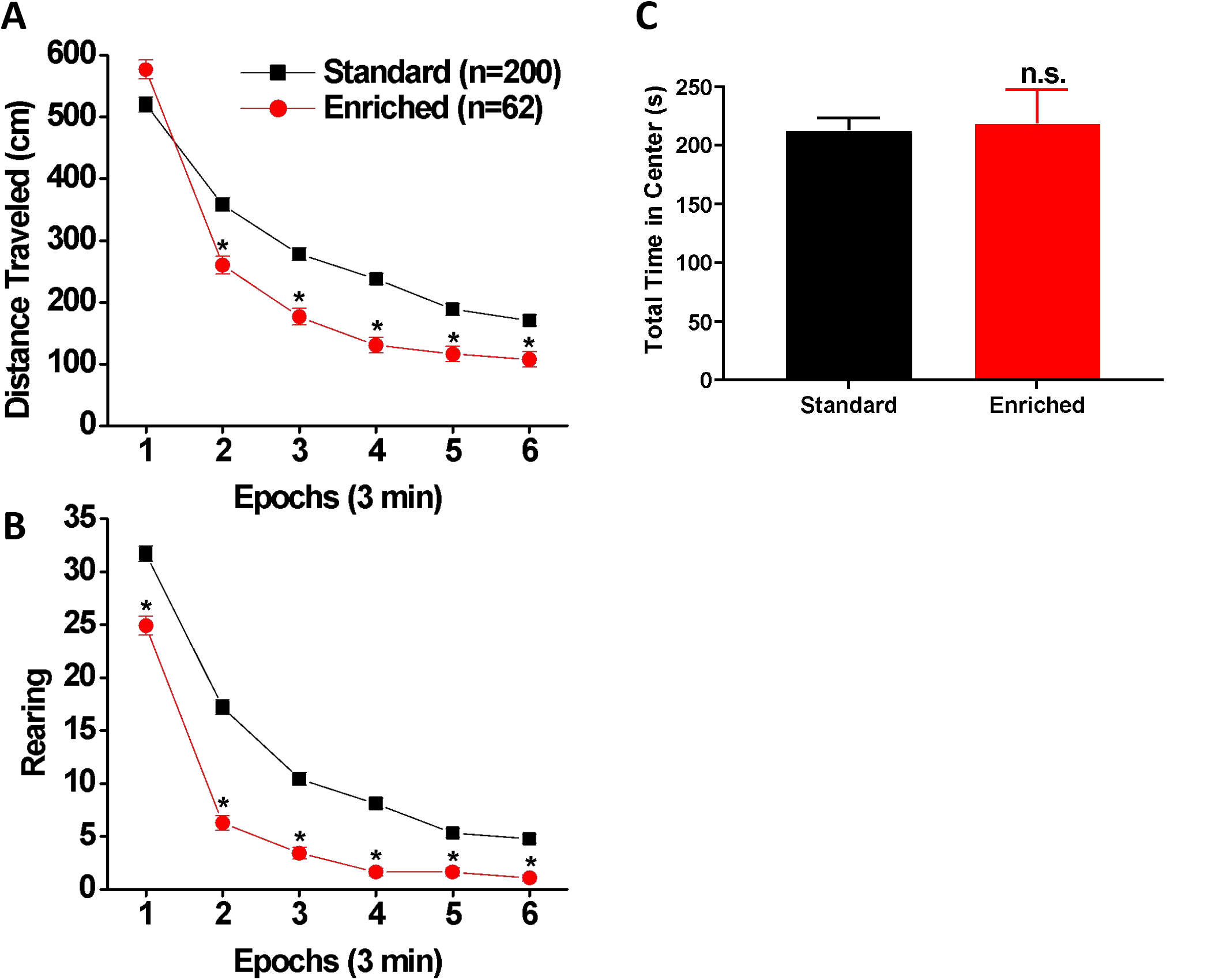
Locomotor response to novelty test. (A and B) Within-session changes in locomotor activity (A: distance traveled; B: rearing) in SH (black square: n = 200) and EE (red circle: n = 62) rats. Data are expressed as means (± SEM) across six 3-min time epochs during the session. * *p* < 0.0083 (= 0.05/6) vs. SH. (C) Mean (± SEM) total time (in seconds) spent by SH and EE rats in the central area of the locomotor chamber.

### Light reinforcement

Raw data from eight SH rats were accidentally lost during one of the light reinforcement sessions, and these rats were consequently dropped from the analysis (thus N for SH=192 and N for EE=64). Focusing on between-session changes, Fig. 4A shows the average number of responses per session during the pre-exposure and light reinforcement phases. During the pre-exposure phase (left-hand panel of Fig. 4A), the EE rats made fewer responses into either port than the SH rats [main effect of housing: F(1, 254) = 231.02, *p* < 0.001, η_p_^2^ = 0.476], while both groups showed sharp declines in responding across the six sessions (intersession habituation) [main effect of session: Wilk’s Ʌ = 0.40, F(5, 250) = 75.05, *p* < 0.001, η_p_^2^ = 0.600]. Responding was similar for both ports, and there were no main effects of port or interactions with port, which was expected because responses into either port are not reinforced during this phase.

**Figure 4:**
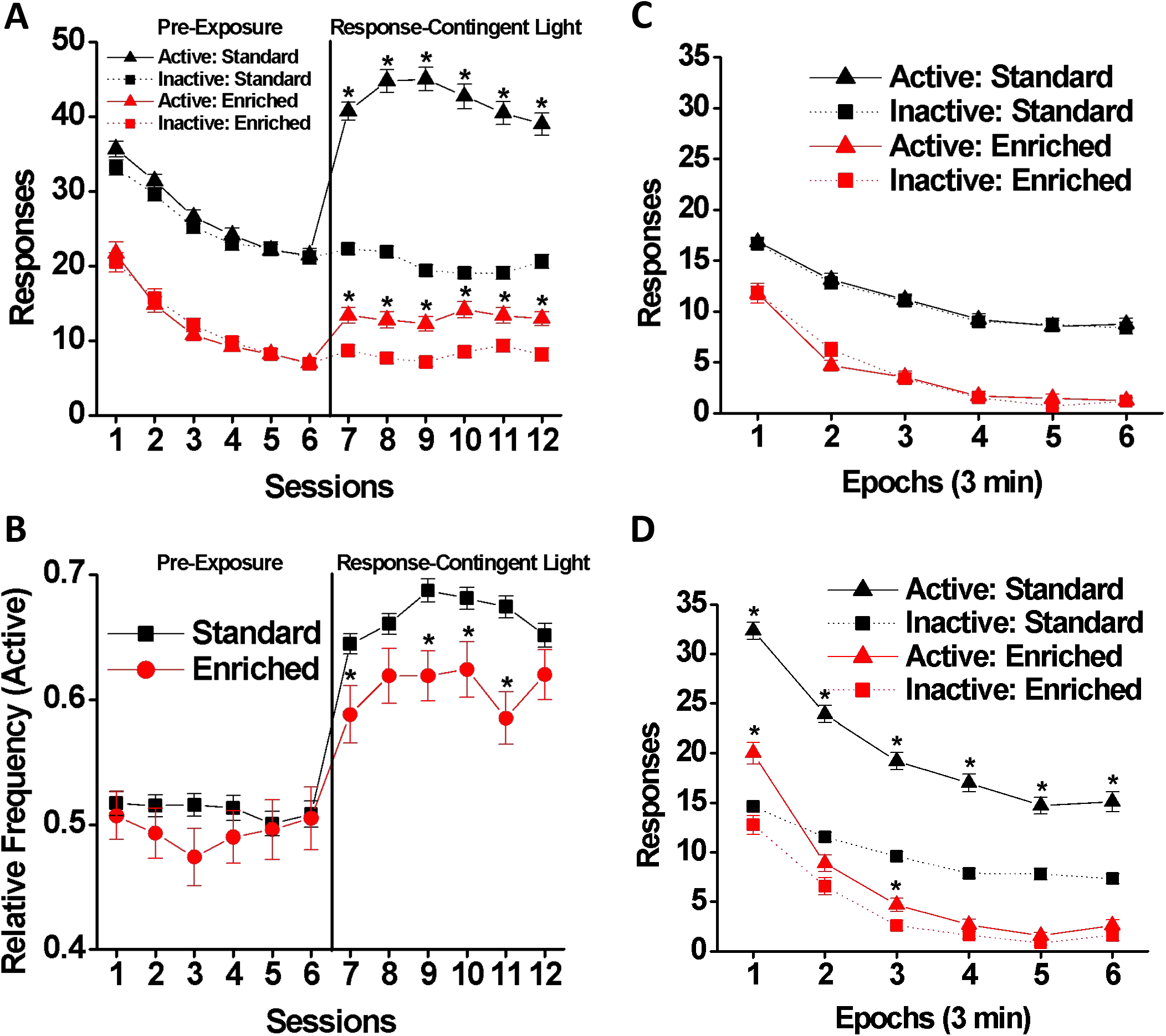
Light reinforcement test. (A and B) Between-session changes in responding during the pre-exposure (left-hand panel) and light reinforcement (right-hand panel) phases of the light reinforcement test by SH (black: n = 200) and EE (red: n = 64) rats. (A) Mean (± SEM) numbers of snout poke responses into the active (triangle) and inactive (square) ports made across six pre-exposure and six light reinforcement sessions. * *p* < 0.0083 (= 0.05/6) vs. inactive port. (B) Mean (± SEM) relative frequency of active responses across six pre-exposure and six light reinforcement sessions. * *p* < 0.0083 vs. SH. (C and D) Within-session changes in responding during (C) pre-exposure and (D) light reinforcement phases. Mean (± SEM) numbers of snout poke responses into the active (triangle) and inactive (square) ports across six time epochs during the session made by the SH (black) and EE (red) rats from the last three sessions. * *p* < 0.0083 vs. inactive port.

When the response-contingent light stimulus was introduced (light reinforcement phase, right-hand panel of Fig. 4A), the number of “active” responses, which produced response-contingent 5-s light onsets, increased significantly relative to “inactive” responding [main effect of port: F(1, 254) = 145.27, *p* < 0.001, η_p_^2^ = 0.364]. Moreover, this increase in active responding relative to inactive responding was smaller for the EE rats than for the SH rats [housing × port interaction: F(1, 254) = 58.10, *p* < 0.001, η_p_^2^ = 0.186]. EE rats made fewer overall responses than the SH rats [main effect for housing: F(1, 254) = 198.18, *p* < 0.001, η_p_^2^ = 0.438], but overall responding did not change significantly across the six sessions [no significant main effect of session: Wilk’s Ʌ = 0.98, F(5, 250) = 1.29, *p* = 0.270]. There were significant two-way housing × session [Wilk’s Ʌ = 0.93, F(5, 250) = 3.82, *p* < 0.01, η_p_^2^ = 0.071] and port × session [Wilk’s Ʌ = 0.93, F(5, 250) = 3.80, *p* < 0.01, η_p_^2^ = 0.71] interactions, and a significant three-way housing × port × session interaction [Wilk’s Ʌ = 0.95, F(5, 250) = 2.66, *p* < 0.05, η_p_^2^ = 0.051] indicated that housing groups had different patterns of responding.

We also looked at the relative frequency of active responding (Fig. 4B). During the pre-exposure phase, the housing groups did not differ in their preference for the “active” port, which also did not change significantly across the six sessions despite overall intersession declines in responding. During the light reinforcement phase, the EE rats’ preference for the light stimulus was weaker than that of the SH rats throughout this phase: main effects of housing [F(1, 254) = 17.16, *p* < 0.001, η_p_^2^ = 0.063] and session [Wilk’s Ʌ = 0.93, F(5, 250) = 3.70, *p* < 0.01] but no significant housing × session interaction [Wilk’s Ʌ = 0.96, F(5, 250) = 2.05, *p* = 0.072]. *Post hoc* comparisons showed that the EE rats had a lower relative frequency of active responding than SH rats in the first, third, fourth, and fifth light reinforcement sessions (*p* < 0.0083).

Figs. 4C and 4D show within-session changes, depicting the mean number of “active” and “inactive” responses across the six 3-min epochs within the session from the last three sessions of the pre-exposure (Fig. 4C) and light reinforcement (Fig. 4D) phases. During pre-exposure, overall responding decreased within the session [main effect of epoch: F (5, 1270) = 136.29, *p* < 0.001, η_p_^2^ = 0.349], but faster in EE rats declined [housing × epoch interaction: F (5, 1270) = 2.81, *p* < 0.05, η_p_^2^ = 0.011]. Overall, the EE rats responded less than the SH rats [main effect of housing: F(1, 254) = 183.77, *p* < 0.001, η_p_^2^ = 0.420], but there was no difference between the number of responses to the “active” versus the “inactive” port. During the light reinforcement phase (Fig. 4D), both groups responded more to the “active” port than to the “inactive” port [main effect of port: F(1, 254) = 108.17, *p* < 0.001, η_p_^2^ = 0.299], but the EE rats made fewer overall responses [main effect of housing: F(1, 254) = 142.63, *p* < 0.001, η_p_^2^ = 0.360], and the difference between active and inactive responding was smaller for the EE rats [housing × port interaction: F(1, 254) = 42.75, *p* < 0.001, η_p_^2^ = 0.144]. Responding by the rats in each housing condition altered at different rates across the epochs [housing × epoch interaction: Wilk’s Ʌ = 0.94, F(5, 250) = 3.23, *p* < 0.01, η_p_^2^ = 0.061]. No significant three-way housing × port × epoch interaction was found. Taken together, the results of the light reinforcement test demonstrated that, compared to the SH rats, the EE rats made fewer responses overall throughout the experiment, and that, when the

response-contingent light was made available, the EE rats showed a reduced tendency to seek light reinforcement.

### Reaction time

As shown in Fig. 5A, the EE rats had a significantly shorter mean reaction time than the SH rats [t(262) = 3.46, p < 0.001, Cohen’s *d* = 0.496]. Fig. 5B shows the average number of trials completed across six time epochs from the last three sessions. There was a significant housing x time epoch interaction [Wilk’s Ʌ = 0.819, F(5, 258) = 11.44, *p* < 0.001, η_p_^2^ = 0.181] as well as significant main effects of housing [F(1, 262) = 119.77, *p* < 0.001, η_p_^2^ = 0.314] and epoch [Wilk’s Ʌ = 0.33, F(5, 258) = 104.34, *p* < 0.001, η_p_^2^ = 0.669]. *Post hoc* comparisons indicated that the EE rats completed more trials in all the six epochs than the SH rats (*p* < 0.0083). Moreover, the EE rats made significantly fewer omissions [t(169.7) = 5.25, *p* < 0.0001, Glass’s *δ* = 0.881] (Fig. 5C) and false alarms per opportunity than the SH rats [t(241.3) = 3.60, *p* < 0.01, Glass’s *δ* = 0.744] (Fig. 5D). Taken together, the results of the reaction time test indicate that the EE rats displayed a better ability to sustain effortful attention and exercised better inhibitory control relative to the SH rats.

**Figure 5:**
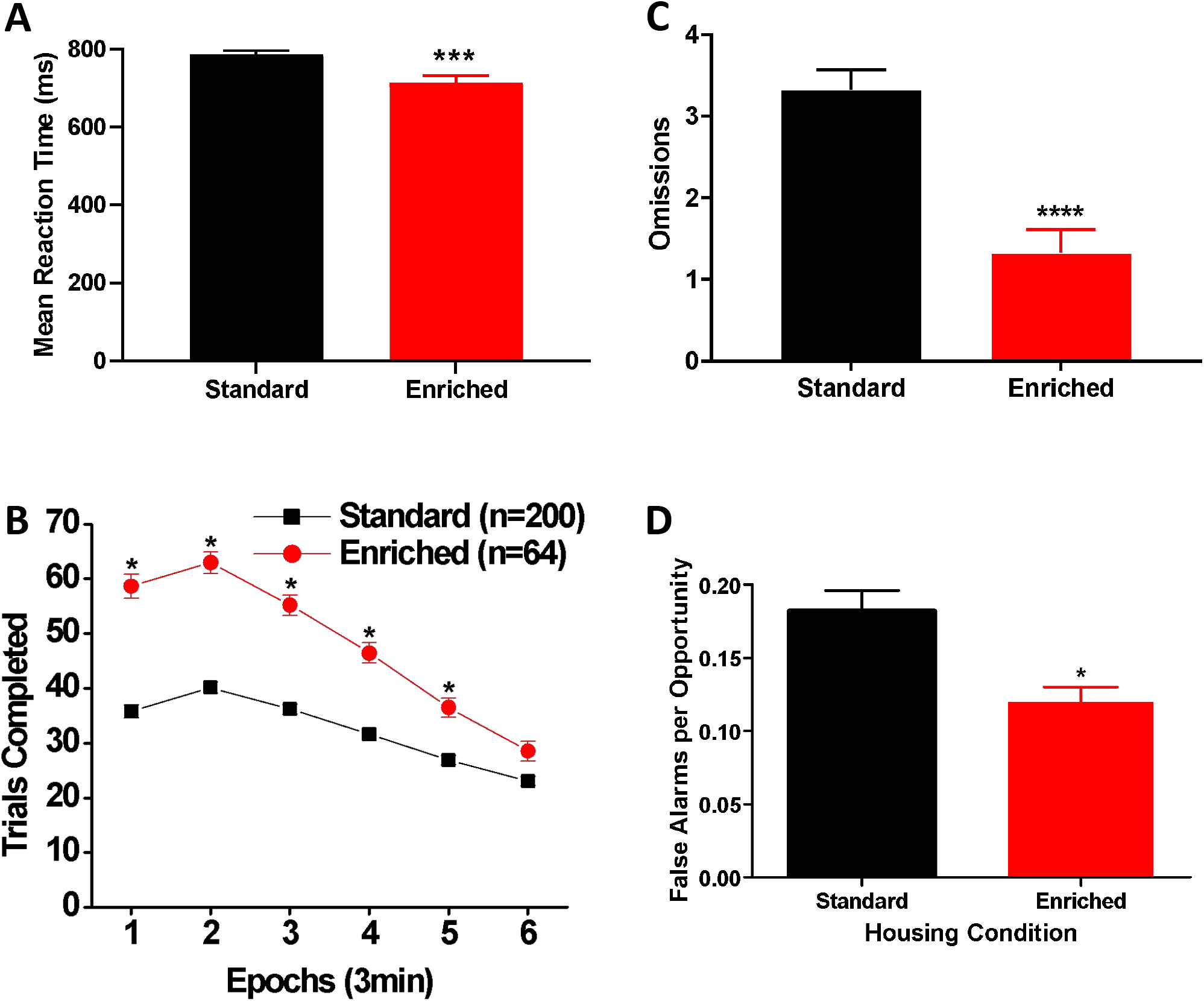
Reaction time test. (A) Mean (± SEM) reaction time (in milliseconds) for SH (black: n=200) and EE (red: n=64) rats. *** *p* < 0.001. (B) Within-session performance by SH and EE rats. Numbers of trails completed by SH and EE rats from the last three sessions are shown. Data are expressed as means (± SEM) across six time epochs during the session. * *p* < 0.0083 (= 0.05/6) vs. SH. (C) Mean (± SEM) number of omissions made by SH and EE rats. **** *p* < 0.0001. (D) Mean (± SEM) number of false alarms per opportunity for SH and EE rats. * *p* < 0.05.

### Patch Depletion

The indifference point functions, showing the reward size when a patch was abandoned as a function of delay to entering a new patch, are depicted in Fig. 6B. As can be seen, as the delay became longer, the switch/indifference points for both groups declined [main effect of delay: Wilk’s Ʌ = 0.09, F(4, 259) = 698.11, *p* < 0.001, η_p_^2^ = 0.915]. The EE rats had lower indifference points than the SH rats, indicating that they earned more reinforcers from the depleting patch prior to switching [main effect of housing: F(1, 262) = 79.17, p < 0.001, η_p_^2^ = 0.232]. There was a significant housing × delay interaction [Wilk’s Ʌ = 0.94, F(4, 259) = 4.18, p < 0.01, η_p_^2^ = 0.061] with the EE rats delaying leaving patches longer than the SH rats. In other words, EE rats discounted the delay incurred when switching to a new patch more than the SH rats. This was more apparent when data were expressed as proportions of the indifference point at 0-s delay (Fig. 6C).

**Figure 6:**
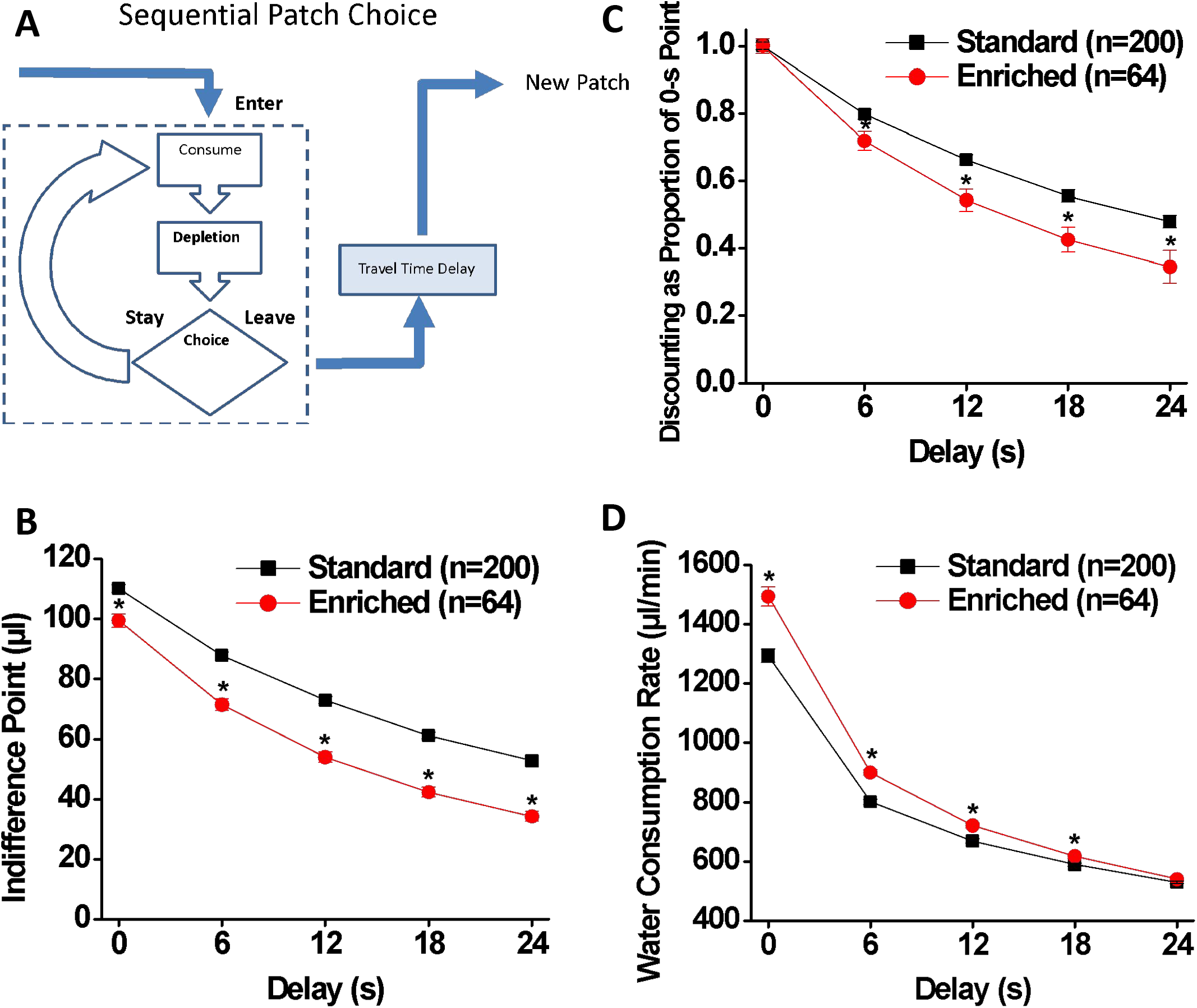
Patch depletion test. (A) Sequential patch depletion procedure. The rats choose between staying in the depleting patch or leaving and moving to a new (undepleted) patch. Longer stays in the depleting patch produce lower rates of immediate water consumption. However, leaving the depleting patch requires enduring a travel time delay, during which the rate of immediate water consumption is zero. Foraging theory predicts that, when confronted with long delays, animals will deplete patches to lower levels before leaving. (B) Indifference points at five delays for SH (black: n = 200) and EE (red: n = 64) rats. The rate of the decrease in the indifference points as function of delay indicates discount rate. (C) Indifference points expressed as a proportion of that at the 0-s delay for SH and EE rats. (D) Water consumption rates at five delays for SH and EE rats. Data are expressed as means ± SEM. * *p* < 0.01 (= 0.05/5) vs. SH.

Steeper discounting in traditional delay discounting paradigms is often viewed as sub-optimal ^3,48^. However, the water consumption rate data suggest that this was not the case (Fig. 6D). There was a significant delay × housing interaction [Wilk’s Ʌ = 0.85, F (4, 259) = 11.11, *p* < 0.001, η_p_^2^ = 0.146] as well as significant main effects of both delay [Wilk’s Ʌ = 0.09, F (4, 259) = 635.02, *p* < 0.001, η_p_^2^ = 0.907] and housing [F (1, 262) = 25.52, *p* < 0.001η_p_^2^ = 0.089]. *Post hoc* comparisons revealed that the EE rats had a higher rate of water intake than the SH rats at 0, 6, 12, and 18-s delays (*p* < 0.01). Thus, the present results indicated that EE rats persisted in patches longer than the SH rats, and that this strategy resulted in higher rates of water consumption and can therefore be viewed as more optimally adjusting to the contingencies imposed by this procedure.

### Pavlovian conditioned approach and conditioned reinforcement

Pavlovian conditioned approach measures the attribution of incentive salience to reward cues, as measured by approach to the cue (sign-tracking), relative to approach to the reward delivery location (goal-tracking). Overall, EE increased sign-tracking compared to SH (Fig. 7). Specifically, EE rats had more lever contacts (the main indicator of sign-tracking behavior; Fig 7A) than SH rats [main effect of housing on lever contacts: F(1, 262) = 31.84, *p* < 0.001, η_p_^2^ = 0.108]. The number of lever contacts increased across sessions [main effect of session: Wilk’s Ʌ = 0.65, F(4, 259) = 34.40, *p* < 0.001, η_p_^2^ = 0.347] but at different rates for the housing groups [housing × session interaction: Wilk’s Ʌ = 0.90, F(4, 259) = 7.54, *p* < 0.001, η_p_^2^ = 0.104]; although *post-hoc* analysis indicated that the EE rats had a larger number of lever contacts than the SH rats in all testing sessions (*p* < 0.01). Fig. 7B shows the number of food cup entries during the presentation of the lever CS, which is indicative of goal-tracking behavior. As can be seen in the figure, the EE rats initially had a higher number of food cup entries than the SH rats and maintained the same level across the sessions, while the initially low number of food cup entries made by the SH rats increased across the sessions and reached the same level as the EE rats by Session 5. However, while approaching significance [F(1, 262) = 3.85, *p* = 0.051], the main effect of housing was not significant. There was also no significant main effect of session [Wilk’s Ʌ = 0.97, F(4, 259) = 2.14, *p* = 0.076] nor a significant housing × session interaction [Wilk’s Ʌ = 0.97, F (4, 259) = 1.84, *p* = 0.121] on food cup entries during lever presentation. However, separate multiple comparisons indicated significant differences between the EE and SH groups in food cup entries during lever presentation on Sessions 1 and 2 (*p* < 0.01). The number of entries into the food cup during inter-trial intervals is shown in Fig. 7C. Due to missing data points caused by system malfunction, two rats in the EE group were dropped from the analysis. There were significant main effects of housing [F(1, 260) = 34.11, *p* < 0.001, η_p_^2^ = 0.116] and session [Wilk’s Ʌ = 0.91, F(4, 257) = 6.15, *p* < 0.001, η_p_^2^ = 0.087] as well as a significant housing × session interaction [Wilk’s Ʌ = 0.91, F (4, 257) = 6.70, *p* < 0.001, η_p_^2^ = 0.094] on food cup entries. *Post-hoc* analysis indicated that EE rats had more inter-trial interval food cup entries than SH rats on Sessions 1 - 4 (*p* < 0.01) but no session 5. In sum, EE increased goal-tracking initially, but biased rats towards sign-tracking in later sessions.

**Figure 7:**
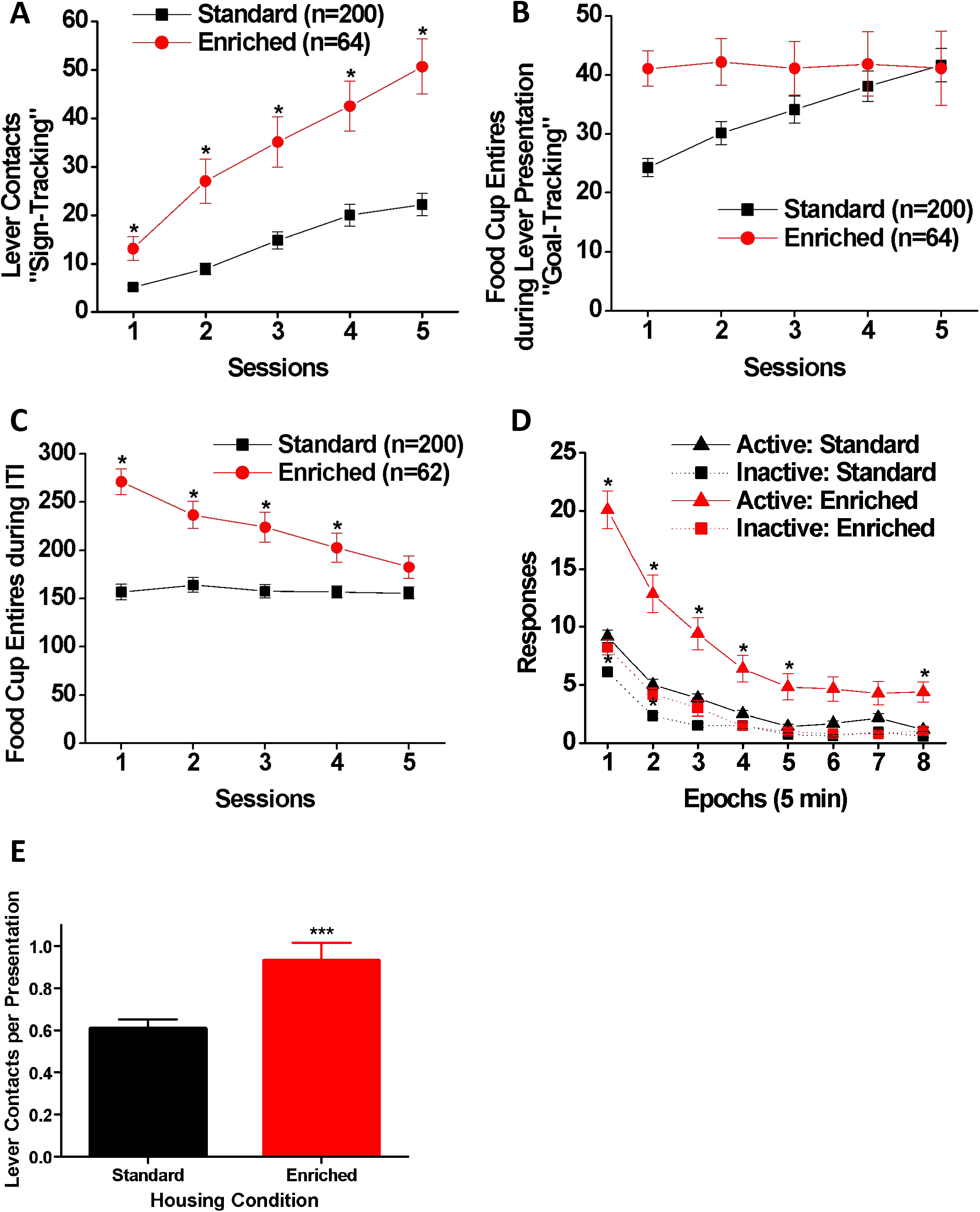
Pavlovian conditioned approach and conditioned reinforcement tests. Numbers of (A) lever contacts and (B) food cup entries made by SH (black: n = 200) and EE (red: n = 64) rats during the presentation of the lever during the Pavlovian conditioned approach test. (C) The number of food-cup entries during the inter-trial interval made by SH (black: n = 200) and EE (red: n = 62) rats. Data are expressed as means (± SEM) across five daily sessions. * *p* < 0.01 (= 0.05/5) vs. SH. (D) Active (triangle) and inactive (square) snout poke responses made by SH (black: n = 200) and EE (red: n = 64) rats during the conditioned reinforcement test. Data are expressed as means (± SEM) across eight time epochs during the session. * *p* < 0.00625 (= 0.05/8) vs. SH. (E) The mean (± SEM) number of lever contacts per presentation for SH and EE rats. *** *p* < 0.001.

Another measure of incentive salience attribution is whether the cue can reinforce behavior even in the absence of the reward, which can be measured during the conditioned reinforcement test. Fig. 7D shows active and inactive responses during the conditioned reinforcement test following the Pavlovian conditioned approach task. The analysis of active snout-poke entries indicated main effects of housing [F(1, 262) = 82.47, *p* < 0.001, η_p_^2^ = 0.239] and epoch [Wilk’s Ʌ = 0.40, F (7, 256) = 54.09, *p* < 0.001, η_p_^2^ = 0.597] as well as a significant housing x epoch interaction [Wilk’s Ʌ = 0.84, F (7, 256) = 6.79, *p* < 0.001, η_p_^2^ = 0.157]. *Post-hoc* analysis indicated that the EE rats had a significantly larger number of active snout-pokes than the SH rats during Epochs 1-5 and 8 (*p* < 0.00625). EE rats also made more inactive snout-pokes as revealed by significant main effects of housing [F(1, 262) = 13.47, *p* < 0.001, η_p_^2^ = 0.049] and epoch [Wilk’s Ʌ = 0.36, F(7, 256) = 66.52, *p* < 0.001, η_p_^2^ = 0.645] as well as a significant housing × bin interaction [Wilk’s Ʌ = 0.91, F(7, 256) = 3.57, *p* < 0.01, η_p_^2^ = 0.089]. However, *post-hoc* analysis indicated a significant group difference during Epochs 1-2 only (*p* < 0.00625), presumably due to floor effects. Moreover, as shown in Fig. 7E, the EE group displayed a higher rate of lever-directed responding compared to the SH group [t(262) = 3.57, *p* < 0.001, Cohen’s *d* = 0.512].

In summary, the EE rats produced significantly more active snout-pokes resulting in the presentation of the lever cue that had been paired with food during the Pavlovian conditioned approach task, and the SH rats.

### Cocaine conditioned cue preference

Overall, EE reduced locomotor activity generally but had no effect on the response to cocaine or preference for the cocaine-paired tactile cue (Fig. 8). Specifically The analysis of the time spent on the cocaine-paired floor (Fig. 8A) showed significant main effects of housing [F(1, 262) = 8.55, *p* < 0.01] and test (pre- and post-test) [F(1, 262) = 105.41, *p* < 0.001] on this measure but no significant housing × test interaction [F(1, 262) = 0.18, *p* = 0.672]. *Post-hoc* tests revealed that both EE and SH groups significantly increased their time spent on the cocaine-paired floor during the post-test compared to the pre-test (*p* < 0.001), suggesting cocaine preference. The EE rats spent significantly more time on the floor to be paired with cocaine than the SH rats during the pre-test (*p* < 0.025), but this difference was not maintained reliably during the post-test (*p* = 0.063), suggesting a lack of group difference in susceptibility to cocaine-cue conditioning. This was underscored by an analysis indicating changes (increases) in the time spent on the cocaine-paired floor between the pre- and post-test did not differ between groups [t(262) = 0.42, *p* = 0.672] (Fig. 8A inset).

**Figure 8:**
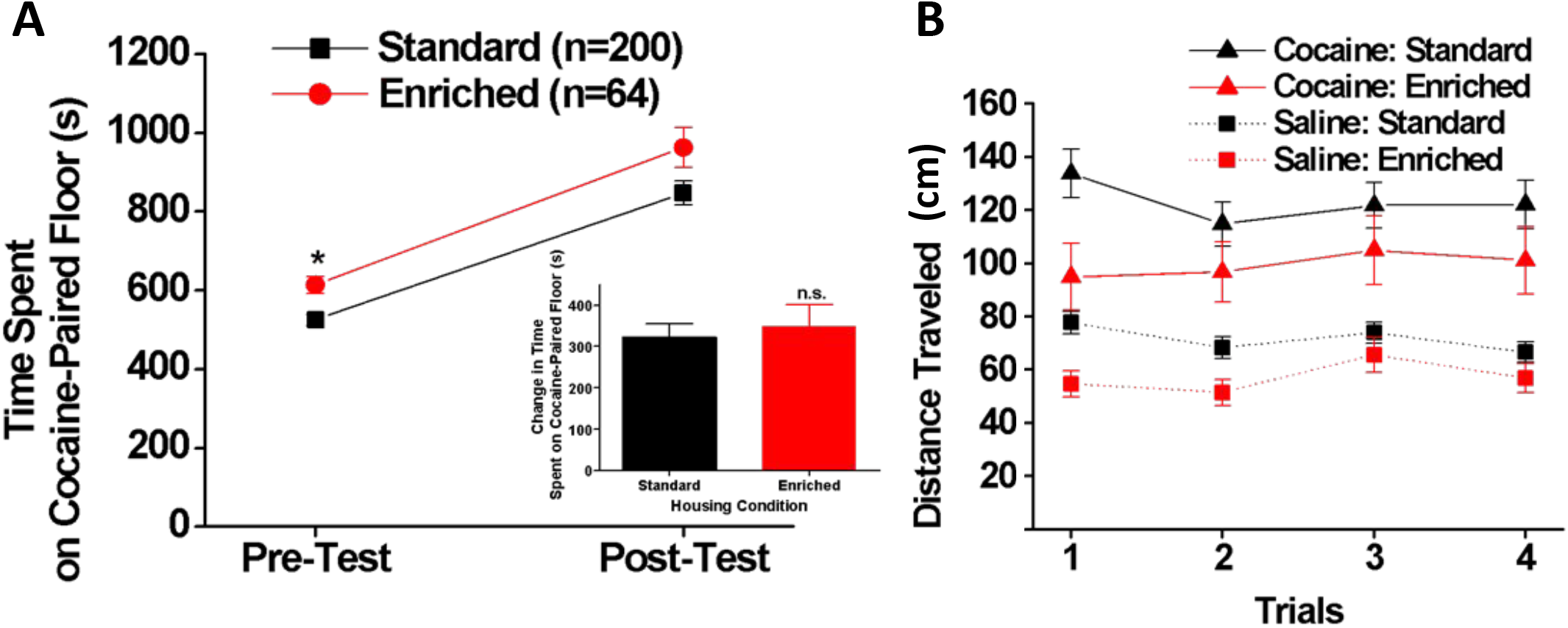
Conditioned cue preference test. (A) Time spent on the cocaine-paired floor by SH (black: n = 200) and EE (red: n = 64) rats on pre- and post-tests. Inset: Change in time spent on the cocaine-paired floor between the two tests. (B) Distance traveled (mm) by SH (black) and EE (red) rats during the conditioning trials with saline (square) and 10 mg/kg cocaine (triangle). Data are expressed as means (± SEM) across four trials.

Fig. 8B shows locomotor activity (distance traveled) during the conditioning trials. A three-way ANOVA (housing, cocaine/saline, trial) revealed a significant main effect of cocaine [F (1, 262) = 108.46, *p* < 0.001], indicating that 10 mg/kg cocaine increased locomotor activity in both EE and SH rats. However, there were no significant main effects of housing condition [F(1, 262) = 3.41, *p* = 0.066] or trial [Wilk’s Ʌ = 0.97, F(3, 260) = 2.45, *p* = 0.064], although these effects approached significance. All two-way and three-way interactions were not significant [housing x cocaine x trial: Wilk’s Ʌ = 0.99, F(3, 260) = 0.49, *p* = 0.690; housing × cocaine: F(1, 262) = 1.05, *p* = 0.308; housing × trial: Wilk’s Ʌ = 0.98, F(3, 260) = 1.63, *p* = 0.182; cocaine × trial: Wilk’s Ʌ = 0.997, F(3, 260) = 0.29, p = 0.834]. In sum, EE had no specific effects on the response to cocaine in this study.

## Discussion

Behavioral regulation is the process by which individuals use their cognitive and emotional resources to control their behavior, thoughts, and emotions to better achieve their goals. It involves the ability to monitor and adjust behavior based on feedback from the environment, as well as the ability to resist distractions and impulses that may interfere with achieving their goals. In our study, these goals included obtaining food and fluid rewards, and learning about the relationship between environmental stimuli and the availability of ingestive and drug rewards. We found that environmental enrichment affected behavior in all of these behavioral paradigms with one notable exception – the response to cocaine. That is, environmentally-enriched rats displayed superior abilities to filter out irrelevant stimuli and focus on relevant stimuli compared to standard-housed rats. This resulted in more efficient performance and was consistent with better regulation (Table 1).

The enrichment in our study included additional sensory, motor, cognitive, and social stimulation elements, as well as multiple resource locations where rats could obtain food and water. Although not tested directly, this range of enriching elements potentially provided multiple, differential stimulus-specific opportunities for subjects to learn stimulus processing and filtering beyond the opportunities provided in standard housing. This raises the possibility that inconsistencies in effects of EE reported in the literature may be attributable to the different enrichment stimulus characteristics provided and the resulting differential learning experiences ^e.g, 49^. This suggestion could be examined experimentally.

Recognition that our enrichment environment included multiple stimulus elements, coupled with our findings, caused us to re-conceptualize and categorize our tests based on the nature of the stimuli presented and the response requirements. This resulted in four *post hoc* categories.

1. *Stimulus Reactivity Tests* included unconditioned or operant responses to a novel environment or low-value reinforcer (Locomotor Response to Novelty, Light Reinforcement). EE rats exhibited relative decreases in Stimulus Reactivity tests.
2. *Adaptive Foraging Tests* included conditioned or operant responses to motivationally high-value reinforcers (reaction time, Patch Depletion, Pavlovian Conditioned Approach). EE rats exhibited relative increases in Adaptive Foraging tests.
3. *Cocaine Sensitivity Tests* consisted of unconditioned and conditioned responses to cocaine’s locomotor and conditioned reinforcing effects (Conditioned Cue Preference). EE rats did not differ from SH rats.
4. *Sociability* was measured in the social reinforcement test, and is placed in a separate category because of the uniqueness of this reinforcer (see below). EE rats exhibited relative increases in Sociability.

By placing our results in the context of previously published literature below, we expand on these ideas before pointing out the study limitations and the avenues for future studies.

### Stimulus Reactivity

One key feature of the EE cages was the presence of interactive toys. This exposure to multiple kinds of visual and tactile stimuli may have reduced rats’ initial response-to-novelty for low-impact stimuli, i.e. stimuli not associated with danger or heightened reward, and may have facilitated subsequent habituation, through a combination of pre-exposure and generalization processes. We contend that our tasks examining response to novelty and light reinforcement exposed subjects to low-impact stimuli, resulting in EE rats exhibiting lower levels of responding and enhanced rates of habituation in these tests. Similar response-to-novelty results were reported for outbred rats ^e.g. 50–53^. Further, our results are consistent with reduced operant responding for light stimuli and faster habituation of the reinforcer effectiveness of light stimuli for EE rats relative to controls reared in standard cages or isolation ^54, 55^.

### Adaptive Foraging

In contrast, the three tasks within the foraging category have the commonality of a motivationally high-value reinforcer. Specifically, rats were reinforced with water in a water-deprived state or learned that a cue predicted a highly-preferred banana-flavored food pellet. Generally, EE rats performed more efficiently, earning more reinforcers in the reaction time and patch depletion tests, and learning the Pavlovian cue-food relationship more rapidly, compared to the SH rats. Further, during the reaction time test, EE rats made fewer premature responses (lower ‘action impulsivity’), thereby displaying a superior ability to attend to task-relevant stimuli.

At first glance, EE rats appeared to have increased ‘choice impulsivity’ compared to the SH rats, because they preferred a sooner, smaller reward compared with SH rats. Others have also reported an increased preference for sooner, smaller rewards over delayed, larger ones by EE rats compared to SH and isolation-reared rats ^e.g. 56^. However, the present study used a sequential patch depletion (i.e., foraging) procedure, which requires the animal to make sequential choices between staying in a “patch” and leaving it for a new patch, rather than typical delay discounting procedures in which the animal has to make mutually exclusive choices between two simultaneously presented discrete alternatives ^32, 42, 57^. One consequence of the parameters in this task (no inter-trial-interval) is that relatively delayed patch-leaving yielded more water reward during sessions. Thus, from an evolutionary and ecological perspective, a preference for sooner, smaller rewards is not necessarily impulsive and can be adaptive, for example, in a natural patchy environment, where resources are scarce and the future is uncertain. In this case, EE rats showed more efficient behavior in the patch depletion task, indicating that their strategy allowed them to obtain more rewards.

In the Pavlovian conditioned approach test, during which the number of rewards is fixed regardless of rats’ responses, EE rats showed enhanced learning rates for Pavlovian conditioned approach compared to SH rats. Remarkably, the EE rats reached asymptotic goal-tracking levels within a single session. SH rats reached this level only after 4-5 days of training. EE rats maintained their high level of goal-tracking even as rates of sign-tracking increased. These observations generally paralleled those from the conditioned reinforcement task, in which the EE rats produced significantly more active snout-pokes producing the food-paired cue than SH rats. In contrast, an earlier study with male Sprague Dawley rats reported higher goal-tracking but lower sign-tracking by EE rats ^58^; perhaps reflective of strain and EE methodological differences. However, like the current study, they reported an increase in bias toward goal-tracking over sessions. That EE promotes goal-tracking in addition to sign-tracking may reflect the EE’s enhancement of responsivity to high-value reward-cues.

### Cocaine Sensitivity

Given our contention that EE increases responsivity to high-value reward-cues, it is a surprise that EE and SH rats did not differ in the time spent on the floor type associated with cocaine (conditioned cue preference (CCP) for cocaine). This could suggest that the above-mentioned link between Pavlovian conditioned approach and reward-cue responsivity does not extend to preference for a cocaine-associated tactile cue. Further, it may indicate that cocaine cue preference and Pavlovian conditioned approach behaviors are driven by different underlying mechanisms. This supposition is supported by our previous work using a large sample of HS rats ^29^ but is not consistent with a previous study showing a correlation between sign-tracking and cue-preference in Sprague-Dawley rats ^47^. This discrepancy is likely due to the differences in genetic structure in the two different outbred rat strains used in these studies ^59, 60^. Strain and methodological differences may also account for the mixed evidence for an effect of EE on cocaine-induced conditioned place preference ^25, 61–66^.

### Sociability

In the social reinforcement task, overall responding was lower in EE rats, but when it occurred, was for a longer duration. This is consistent with numerous studies reporting increases in social behavior in EE animals relative to non-EE controls, especially social interaction and exploration ^e.g. 67–70^, although some studies have not observed this effect ^e.g. 71–73^. This inconsistency could be due to methodological differences in the enrichment protocols, including the number of animals per cage, cage size, types of enrichment, sex and strain of animals, age of animals at the onset of EE, duration of EE, and types of controls employed. As we have suggested, different EE components may influence different aspects of behavior, and data from studies enriching the physical environment has been shown to increase social play behavior in adolescent male rats relative to SH controls (Morley-Fletcher *et al.*, 2003).

### Study limitations

Our basic premise when interpreting the data was that the observed effects were due to interactions with specific aspects of the enriched environment. Unfortunately, additional parametric studies would be required to identify which aspects mediated the effects nor whether there are “dose” effects for some or all EE aspects. Another limitation is that the present study included only male rats. EE may affect behavioral regulation in males and females differently depending on the mediating factors such as stress reactivity, and sex differences have been reported in some of the psychological traits that were examined in the present study. We focused on only one sex in part to avoid co-housing males and females in the same enriched environment. However, future studies to examine the effect on enriched environments on females are clearly needed. Yet another limitation was the fixed order of testing in our extensive behavioral phenotyping regime, which may have resulted in carryover effects, and certainly led to rats undergoing tests at different ages. For example, locomotor response to novelty was examined when rats were still in adolescence, but rats were tested on later tasks like patch depletion in adulthood. Some of the traits studied in the present study have been shown to change with age, for example, sensation seeking/novelty seeking phenotypes ^74^, and interactions between enrichment and age on these assays are unknown. Accordingly, age controls would be recommended in future studies of the effects of EE on behavioral regulation. Finally, while we assume that the enriched environment led to enduring changes at the molecular, cellular and circuit levels, we have not yet attempted to understand the changes that mediate these behavioral effects.

## Conclusions: Implications for animal behavioral studies

One clear implication of the present study is that housing conditions can greatly influence results obtained from behavioral tests that use rats as subjects. Presumably these effects could alter the effects of pharmacological or genetic manipulations that are frequently used in conjunction with behavioral measurements such as the ones examined in this present study. Currently, most behavioral studies using rats as subjects are undertaken with animals housed in so-called “standard” laboratory housing, which, with its limited environmental and social complexity, provides relatively little sensory, motor, cognitive, and social stimulation. It is therefore plausible that keeping laboratory animals in such housing conditions may systematically influence brain development and functioning in ways that compromise their ability to adapt to environmental challenges. If true, this can potentially have tremendous impacts on the outcome of behavioral studies. Indeed, significant gene × environment interactions have been observed on behavioral measures in a number of rodent models of human disorders, indicating that behavioral phenotypes of animals with identical genotypes can differ depending on housing conditions ^75, 76^. Thus, researchers must take into consideration these possible consequences of keeping experimental animals in “standard” conditions if they are serious about the generalizability of findings of animal research investigating the mechanisms underlying human disorders. In particular, it warrants serious consideration whether “enrichment” should be considered as a therapeutic intervention or the standard condition required for developing a normal brain ^76, 77^.

## Acknowledgements

The authors would like to thank Brady Thompson, M.S., for his comments on an earlier draft of this manuscript. This work was funded by a center grant from the National Institute of Drug Abuse to AAP, LCSW, DMD, and PJM (DA037844), to SHM (DA046077), and by a grant to PJM from the Office of the Director, National Institutes of Health, and the National Institute on Alcohol Abuse and Alcoholism (AA024112).

## Data Availability

All data generated or analyzed during this study are included in this published article [and its supplementary information files]. This study is reported in accordance with ARRIVE guidelines (https://arriveguidelines.org).

